# Ubiquitylation by the GID/CTLH complex regulates the metabolic and innate immune response of macrophages to infection by *Mycobacterium tuberculosis*

**DOI:** 10.64898/2026.04.24.720540

**Authors:** Nelson V. Simwela, Luana Johnston, Christopher M. Sassetti, David G. Russell

## Abstract

The GID/CTLH E3 ligase complex is implicated in several biological processes, yet its full substrate repertoire remains poorly defined. We recently identified the complex as a broad modulator of macrophage responses to *Mycobacterium tuberculosis* (Mtb) infection. Here, we use label-free proteomics and diGly capture analysis of Mtb-infected macrophages to define the GID/CTLH-dependent ubiquitylome. We identify thousands of dynamically altered ubiquitylation sites, with strong enrichment among proteins involved in cellular metabolism and innate immune signaling. Concurrent proteome analysis revealed extensive rewiring in GID/CTLH-deficient macrophages, with >90% of enriched pathways among increased proteins consisting of metabolic targets. Notably, inhibitory phosphatases (PTEN, INPP5D) also emerged as candidate substrates. Functional studies revealed proteasome-dependent stabilization of PTEN and INPP5D in GID/CTLH-deficient macrophages with each phosphatase individually exerting an influence on Mtb intracellular survival. Together, our study defines a GID/CTLH-dependent ubiquitylome in macrophages and identifies the complex as a central regulator of metabolism and antimicrobial immunity.

**Author summary:** *Mycobacterium tuberculosis* (Mtb), the bacterium that causes tuberculosis (TB), survives and replicates within macrophages, key immune cells that normally eliminate pathogens. How macrophages control their internal cellular environment in response to infection remains incompletely understood. One such important cellular control system is ubiquitylation, in which proteins are tagged with ubiquitin to determine their functional fate or target them for degradation. We recently identified the GID/CTLH E3 ligase ubiquitylation complex as a critical modulator of macrophage antimicrobial responses to Mtb. Here, we used proteomics approaches to define the proteins controlled by the GID/CTLH complex in Mtb-infected macrophages. We found that this complex ubiquitylates a broad network of proteins involved in cellular metabolism and immune signaling. When the complex is disrupted, macrophages undergo extensive metabolic reprogramming, particularly increased mitochondrial energy production, while showing reduced inflammatory signaling. Despite this dampened immune response, these cells are better able to restrict Mtb growth. We also identified the phosphatases PTEN and INPP5D as targets controlled by the GID/CTLH complex that independently influence intracellular bacterial survival. Our findings demonstrate that the GID/CTLH complex is a critical regulator of metabolism and immune function, shaping the outcomes of Mtb infection.

## Introduction

Macrophages are highly plastic immune cells that eliminate pathogens through potent antimicrobial mechanisms while balancing their own survival and maintaining tissue homeostasis. In the context of *Mycobacterium tuberculosis* (Mtb) infection, the causative agent of human tuberculosis (TB), this balance is particularly critical, as optimal antimicrobial responses contain bacterial growth, whereas insufficient responses promote bacterial replication, host cell death, and disease dissemination [1–3]. Understanding the molecular mechanisms that regulate intracellular signaling and antimicrobial pathways in Mtb-infected macrophages can therefore provide important insights for the development of novel strategies to treat or prevent TB. One such mechanism is protein ubiquitylation, a post-translational modification that marks proteins with a ubiquitin tag either for intrinsic cell signaling or degradation by the 26S proteasome. In macrophages, ubiquitylation fine-tunes innate immune responses by controlling the activation, signaling capacity, stability, and turnover of proteins and molecular complexes in response to danger signals [4]. This dynamic regulation is particularly important because, unlike specialized immune cells such as dendritic cells, macrophages are often heterogeneous and must perform multiple functions across the body, ranging from homeostatic activities such as tissue repair and wound healing, to inflammatory activities such as antimicrobial defense and antigen presentation [3, 5]. Moreover, ubiquitylation can directly target intracellular pathogens, including Mtb, for autophagic degradation within phagolysosomes [6–8].

Protein ubiquitylation is mediated by a hierarchical cascade of E1 activating enzymes, E2 conjugating enzymes, and E3 ubiquitin ligases, with E3 ligases conferring substrate specificity and temporal control of individual substrate modification [9]. Among E3 ligases, multi-subunit complexes are particularly versatile, as they recruit diverse substrates through modular scaffolds and adaptor components [10]. One example of such a complex is the glucose-induced degradation (GID) E3 ligase which was originally identified in yeast as a regulator of carbohydrate metabolism [11]. The complex is evolutionarily conserved in mammals as the C-terminal to LisH (CTLH) E3 ubiquitin ligase [12]. The mammalian GID/CTLH E3 ligase complex is composed of the RING-domain catalytic subunits RMND5A and MAEA, together with multiple scaffold and adaptor proteins, including GID8, GID4, WDR26, MKLN1, and YPEL5, that mediate complex stability, substrate recognition, and recruitment [13, 14]. Although some studies have reported interactions of the mammalian GID/CTLH complex with metabolic enzymes such as PKM2 and LDHA [15], including both canonical and non-canonical ubiquitylation of metabolic proteins such HMG-CoA Synthase 1, AMPK and NMNAT1 [16–18], these effects appear largely context-dependent, and the contribution of the complex to metabolic control in mammalian cells remains unclear [14, 19]. In contrast, genetic and biochemical studies have implicated the mammalian GID/CTLH complex in a broad spectrum of cellular processes, including cell proliferation, cell cycle control, antibody maturation, DNA replication, stress responses, and the coordination of proteostatic and signaling networks that support cellular adaptation under both physiological conditions and disease states such as cancer cell plasticity and neurodegeneration [13, 14, 19–24]. In light of the broad range of activities in which the complex is implicated, it is important that the physiological substrates of the complex and its systems-level functions are more fully elucidated.

Through a genome-wide CRISPR-Cas9 knockout screen in Mtb-infected macrophages, we recently discovered that the GID/CTLH complex functions as a negative modulator of macrophage antimicrobial responses [25]. Loss of individual complex subunits enhances macrophage control of Mtb infection, indicating that, under basal and infection-induced conditions, the complex constrains host defense pathways. Transcriptional and functional analyses revealed that GID/CTLH-deficient macrophages activate a wide spectrum of antimicrobial responses, including autophagy and nutritional immunity, despite broadly suppressing canonical cytokine-dependent antimicrobial pathways. However, the molecular mechanisms underlying this phenotype are unknown. In particular, it remains unclear how GID/CTLH-mediated ubiquitylation shapes macrophage signaling networks during infection and which substrates may be linked to intracellular bacteria control.

In this report, we define the substrate landscape and functional impact of the GID/CTLH complex during Mtb infection of macrophages. By combining label-free quantitative (LFQ) proteomics with diGly ubiquitin remnant profiling, we map the GID/CTLH-dependent ubiquitylome and reveal widespread, dynamic changes in protein abundance and ubiquitylation in response to infection and loss of GID/CTLH activity. Our analyses uncover a strong enrichment for metabolic and innate immune targets among GID/CTLH-modulated proteins and demonstrate extensive metabolic rewiring in Mtb-infected GID/CTLH-deficient macrophages. Notably, we identify the inhibitory phosphatases PTEN and INPP5D as proteasome-degraded substrates of the GID/CTLH complex and establish their individual roles in controlling intracellular Mtb survival. Together, these findings implicate the GID/CTLH complex as a central regulator of host cellular pathways that integrate cellular metabolism and antimicrobial immunity, providing new insights into how ubiquitin-mediated signaling fine-tunes host defense during Mtb infection of macrophages.

## Results

### Global and ubiquitin-remnant proteomics to uncover GID/CTLH E3 ligase substrates in Mtb-infected macrophages

To define the role of the GID/CTLH complex in host proteome responses during Mtb infection, we performed LFQ-based global proteomics and ubiquitylomics in infected scramble control, GID8^-/-^, and MAEA^-/-^ Hoxb8 bone marrow-derived macrophages (BMDM^h^s). We previously identified five of the twelve members of the GID/CTLH complex (MAEA, GID8, WDR26, YPEL5, and UBE2H) and experimentally validated up to seven individual members (the five identified hits plus MKLN1 and RANBP9) as determinants of Mtb replication in BMDM^h^s [25]. Functional analyses revealed that antimicrobial responses in macrophages deficient for the examined individual complex members were broadly similar. We therefore focused our proteomics analysis on GID8^-/-^ and MAEA^-/-^ BMDM^h^s to generate complementary datasets and because MAEA forms part of the catalytic domain of the GID/CTLH complex, whereas GID8 is required for structural tethering and integrity of the entire complex [14, 19, 26].

Scramble control, GID8^-/-^, and MAEA^-/-^ BMDM^h^s were infected with wild-type Mtb Erdman strain at a multiplicity of infection (MOI) of 1.5 for 48 hours. Cells were methanol-fixed overnight and lysed, followed by tryptic digestion and di-glycine (diGly) peptide enrichment. Samples were then analyzed to quantify global protein abundance and ubiquitylation dynamics, as summarized in Fig. 1A. Principal component analysis (PCA) of LFQ proteome data revealed clear segregation of scramble, GID8^-/-^, and MAEA^-/-^ samples, with tight clustering of biological replicates (Fig. 1B). The first principal component (PC1; 55.7% variance) separated scramble controls from both knockout conditions, while PC2 (10.4% variance) further distinguished GID8^-/-^ and MAEA^-/-^ BMDM^h^s, indicating both shared and subunit-specific proteomic effects. Across all conditions, 5,087 proteins were identified. Differential protein abundance analysis demonstrated extensive proteome remodeling following disruption of the GID/CTLH complex subunits. Relative to scramble controls, GID8^-/-^ BMDM^h^s exhibited 1,565 increased and 1,216 decreased proteins, whereas MAEA^-/-^ BMDM^h^s showed 1,616 increased and 1,239 decreased proteins (Fig. 1C, S1 Data). These widespread changes indicate that loss of either GID8 or MAEA profoundly alters host protein homeostasis in BMDM^h^s during Mtb infection. Visualization of differential protein abundance using volcano plots (Fig. S1) revealed that GID8 and MAEA were among the most significantly decreased proteins in their respective knockout mutant cell lines, internally validating the robustness of our experimental approach. Notably, several other components of the GID/CTLH complex were consistently decreased in GID8^-/-^ (ARMC8, MAEA, and UBE2H) and MAEA^-/-^ (RMND5A and UBE2H) BMDM^h^s (Fig. S1, S1 Data). This coordinated reduction suggests destabilization of the entire GID/CTLH complex upon loss of individual subunits, as has been previously observed [19], consistent with interdependent maintenance of complex integrity. In contrast, numerous proteins displayed significantly increased abundance in both knockout conditions, including NCOA3, EPHX1, SFXN5, and ABHD13, pointing to shared downstream pathways that may normally be restrained by GID/CTLH-mediated ubiquitylation. Together, these data demonstrate that disruption of either GID8 or MAEA leads to coordinated reductions in the abundance of other GID/CTLH complex subunits and extensive remodeling of the host proteome in BMDM^h^s during Mtb infection.

**Fig 1.**
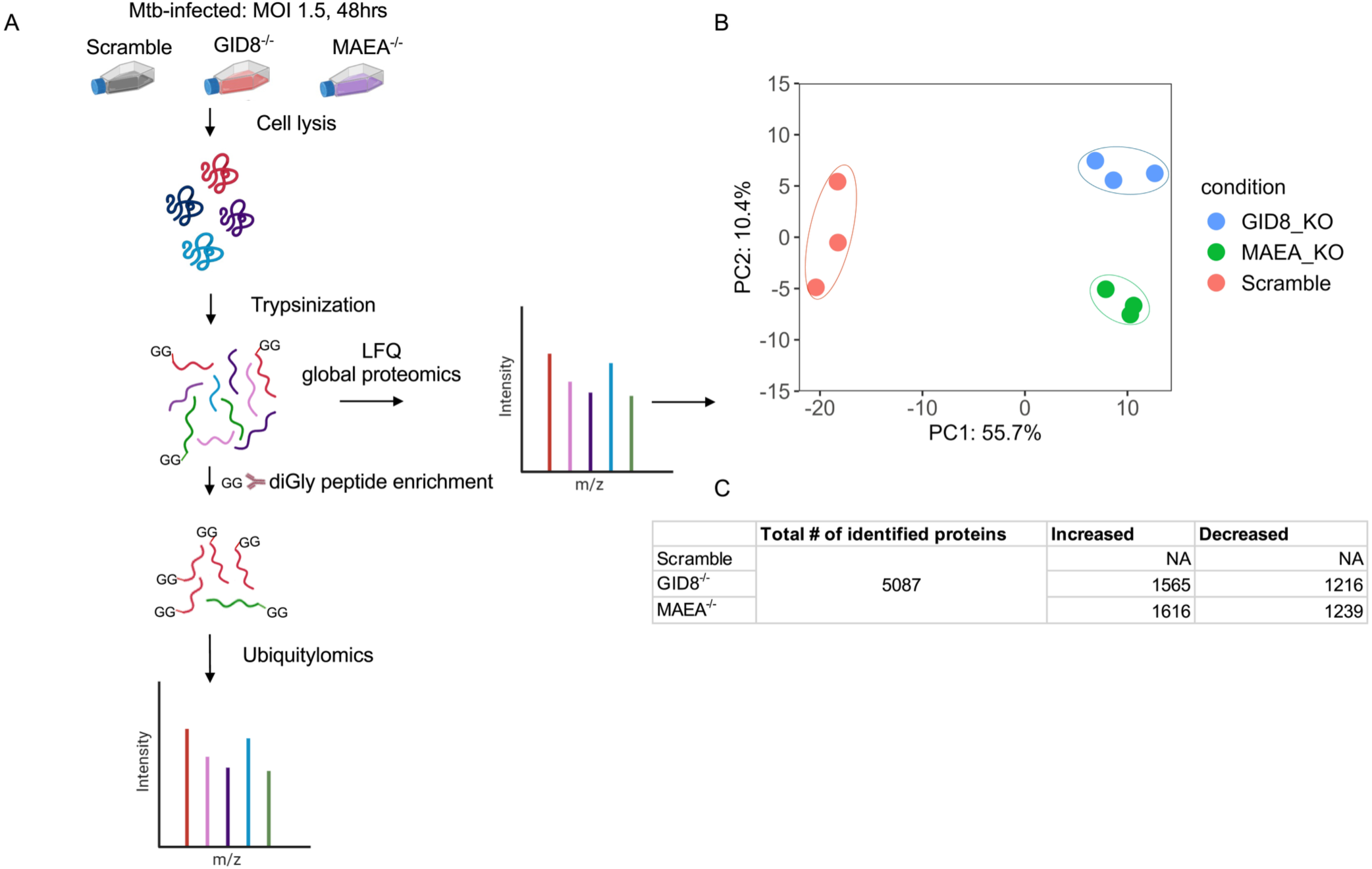
Quantitative label free and diGly proteomics to identify substrates of the GID/CTLH E3 ligase complex in Mtb-infected BMDM^h^s. **A.** Overview of the experimental workflow for label-free and quantitative ubiquitin diGly proteomics. Approximately 1 x 10⁸ Mtb-infected wild-type (scramble control) and GID/CTLH complex-deficient BMDM^h^s (GID8^-/-^ and MAEA^-/-^) were fixed overnight in pre-chilled 90% methanol, lysed, and digested with trypsin. An aliquot of the resulting trypsin digested peptides was subjected to label-free quantification (LFQ) to assess the global proteome. The remaining peptides were enriched for ubiquitylated (diGly-modified) peptides and analyzed to profile global ubiquitylation. Experiments were performed with three biological replicates per genotype. **B.** Principal component analysis (PCA) of LFQ global proteome data showing distinct separation of proteomic responses among scramble control, GID8^-/-^ and MAEA^-/-^ BMDM^h^s. **C.** Illustrated summary table showing the number of identified proteins in the experiment and the differentially abundant proteins in GID8^-/-^ and MAEA^-/-^BMDM^h^s (Data S1). Significant increase or decrease in protein abundance was defined as *p* ≤ 0.05 with log₂ fold change ≥ 0.3 (Up) or ≤ −0.3 (Down).

### The proteomic landscape induced by Mtb infection is highly similar between GID8^-/-^ and MAEA^-/-^macrophages

GID8^-/-^ and MAEA^-/-^ BMDM^h^s showed conserved transcriptional responses following Mtb infection [25] and separated close together in our LFQ proteomic analyses (Fig. 1B). To further quantify the extent of this conservation, we compared proteome-wide changes between the two knockouts. Venn diagram analysis revealed a strong overlap in proteins whose abundance was altered upon loss of GID8 or MAEA (Fig. 2A, S1 Data). Among proteins with increased abundance, 1,398 proteins (78.4%) were commonly increased in both GID8^-/-^ and MAEA^-/-^ BMDM^h^s, whereas 167 proteins (9.4%) and 218 proteins (12.2%) were uniquely increased following GID8 or MAEA knockout, respectively. Similarly, decreased proteins showed substantial concordance, with 992 proteins (67.8%) decreased in both conditions and smaller subsets uniquely reduced upon GID8 (224 proteins, 15.3%) or MAEA (247 proteins, 16.9%) depletion. We also evaluated global proteomic similarities using hierarchical clustering and heatmap analyses of differentially abundant proteins between the two knockout conditions (Fig. 2B). Mtb-infected BMDM^h^ samples segregated by genotype, with GID8^-/-^ and MAEA^-/-^ clustering closer together and remaining clearly distinct from scramble controls (Fig. 2B). Proteins increased upon GID8 or MAEA depletion showed coordinated increases in abundance across both mutants, whereas proteins with reduced abundance showed consistent decreases relative to controls. The overall magnitude and direction of protein abundance changes were highly similar between GID8^-/-^ and MAEA^-/-^ BMDM^h^s. These results demonstrate that loss of GID8 or MAEA induces comparable proteomic alterations, in agreement with our previous transcriptional findings [25], and support a shared or cooperative role for individual members of the complex in maintaining cellular protein homeostasis.

**Fig 2.**
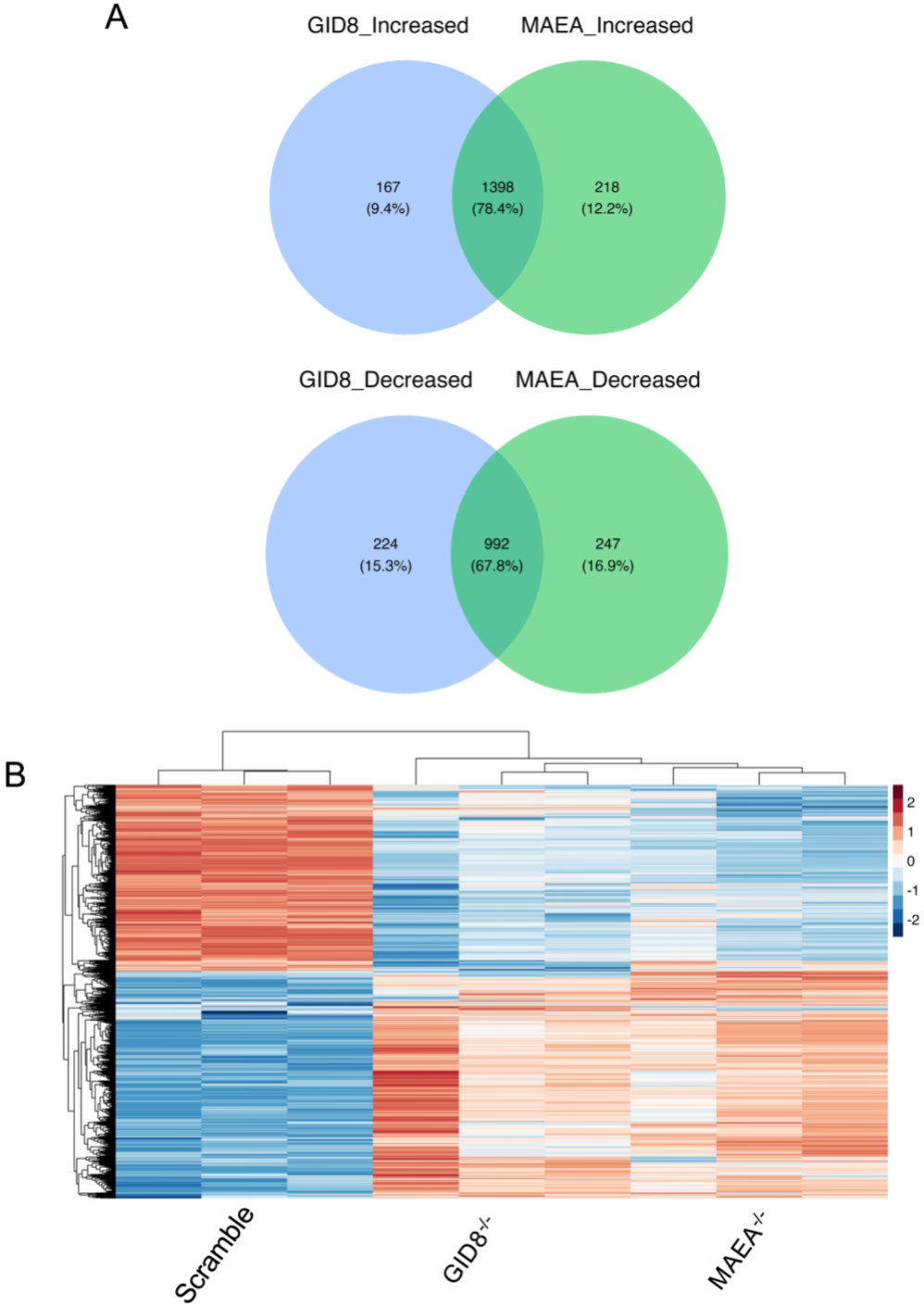
The global proteomic response to Mtb infection is broadly similar in GID8^-/-^ and MAEA^-/-^ BMDM^h^s. **A**. Venn diagrams showing overlaps between increased and decreased differentially abundant proteins in Mtb-infected GID8^-/-^ and MAEA^-/-^ BMDM^h^s, highlighting shared and conserved responses. **B**. Heatmap of all differentially abundant proteins in Mtb-infected GID8^-/-^ and MAEA^-/-^ BMDM^h^s, illustrating similarities and conserved abundance patterns.

### Metabolic processes overwhelmingly dominate the differentially abundant proteome in Mtb-infected GID8^-/-^ and MAEA^-/-^ macrophages

We next performed pathway enrichment analysis [27] on differentially abundant proteins in GID8^-/-^ and MAEA^-/-^ Mtb-infected BMDM^h^s to identify overrepresented biological pathways. Given that both GID8^-/-^and MAEA^-/-^ BMDM^h^s elicited very similar proteomic responses (Fig. 1B, Fig. 2), we focused our pathway enrichment on proteins that were commonly increased or decreased (S1 Data) between the mutants as our input. Enrichment analysis revealed that metabolic processes overwhelmingly dominate the differentially abundant proteome in Mtb*-*infected GID8^-/-^ and MAEA^-/-^ BMDM^h^s. Among the increased proteins, mitochondrial respiration and oxidative phosphorylation emerged as the most significantly enriched functional categories (Fig. 3A, S2 Data). Core pathways related to cellular respiration, aerobic respiration, and electron transport chain activity were enriched, including mitochondrial respiratory chain complex assembly, with prominent enrichment of Complex I (NADH dehydrogenase) assembly, respirasome formation, and ATP synthesis coupled to electron transport. Enrichment of multiple terms associated with proton motive force-driven ATP synthesis and inner mitochondrial membrane organization suggests not only increased respiratory activity but also structural remodeling of mitochondrial cristae to accommodate elevated bioenergetic capacity. Consistent with this, pathways involved in mitochondrial gene expression, mitochondrial translation, rRNA processing, and ribosome biogenesis were increased, indicating coordinated induction of the mitochondrial protein synthesis machinery to sustain enhanced oxidative phosphorylation capacity. These results support a state of significantly enhanced mitochondrial biogenesis and respiratory capacity in GID8^-/-^ and MAEA^-/-^ BMDM^h^s during Mtb infection, in line with our previous functional assays showing that GID/CTLH-deficient BMDM^h^s maintain a state of metabolic resilience upon infection with Mtb [25].

**Fig 3.**
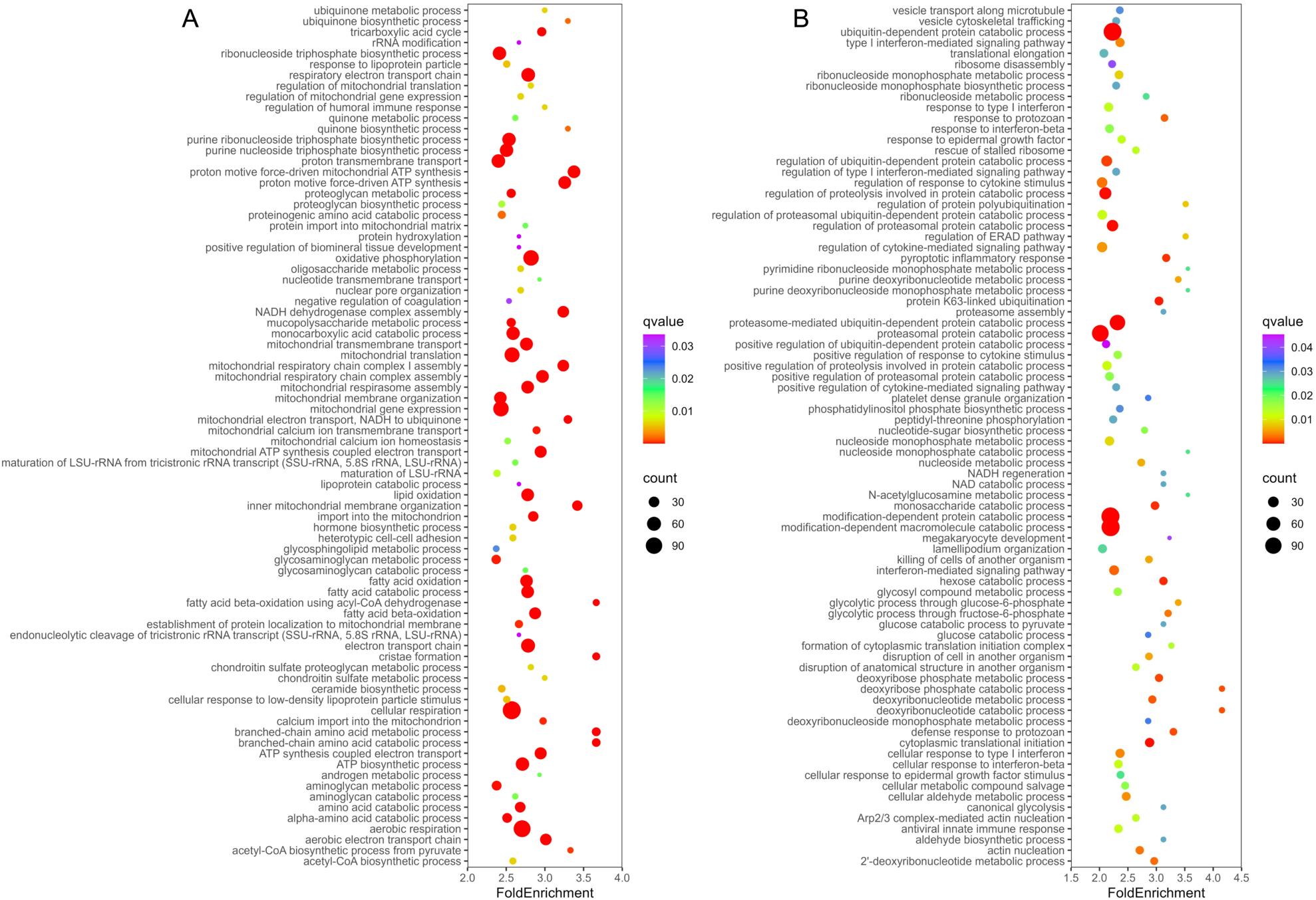
Overrepresented biological pathways among DAPs in Mtb-infected GID8^-/-^ and MAEA^-/-^ BMDM^h^s are dominantly metabolic. **A**. Top enriched pathways among commonly increased proteins in GID8^-/-^ and MAEA^-/-^ BMDM^h^s. **B.** Top enriched pathways among commonly decreased proteins in GID8^-/-^ and MAEA^-/-^BMDM^h^s. Complete datasets are provided in Data S2.

In parallel with enhanced oxidative phosphorylation, lipid and fatty acid catabolic pathways were enriched amongst proteins of increased abundance. These included fatty acid β-oxidation, acyl-CoA metabolic and biosynthetic processes, lipid oxidation, and monocarboxylic and carboxylic acid catabolism. These patterns suggest a metabolic shift toward increased utilization of lipid substrates as a primary fuel source for mitochondrial respiration. Enrichment of the tricarboxylic acid (TCA) cycle and acetyl-CoA biosynthetic pathways further supports enhanced coupling between substrate oxidation and mitochondrial ATP production as has been observed previously in Mtb-infected macrophages [28]. Moreover, increase in organic acid, oxoacid, and amino acid catabolic processes, including branched-chain and α-amino acid catabolism, suggests broader remodeling of intermediary metabolism to provide reducing equivalents (NADH and FADH₂) and metabolic intermediates to support respiratory chain activity. Additional enrichment among increased proteins was observed for pathways involved in mitochondrial transport, protein import into the mitochondrial matrix and inner membrane, and monoatomic ion transmembrane transport, including calcium transport. In macrophages, these processes are essential for maintaining mitochondrial proteome integrity, controlling mitochondrial enzyme activity, and supporting optimal respiratory chain function [29]. The concurrent enrichment of reactive oxygen species (ROS) metabolic pathways suggests altered redox homeostasis accompanying elevated electron transport activity, consistent with increased electron flux through the respiratory chain. This could contribute to the enhanced control of Mtb observed in GID/CTLH knockout BMDM^h^s, in line with our prior findings that Mtb recovered from these macrophages are oxidatively stressed [25].

In contrast, pathway enrichment analysis of proteins of decreased abundance revealed a pronounced overrepresentation of protein quality control and innate immune signaling pathways (Fig. 3B, S2 Data). Processes related to ubiquitin-dependent protein catabolism were strongly enriched, including proteasome-mediated protein degradation, proteasomal protein catabolic processes, ER-associated degradation, and regulation of proteolysis. Multiple ubiquitylation-related pathways, such as protein ubiquitylation, K63-linked ubiquitylation, polyubiquitylation, and proteasome assembly, were also enriched, indicating coordinated attenuation of the ubiquitin-proteasome system. This was accompanied by enrichment of pathways involved in post-translational protein modification, phosphorylation and dephosphorylation, and regulation of protein-containing complex assembly, suggesting broad impairment of proteostasis and signaling protein turnover in Mtb-infected GID8^-/-^ and MAEA^-/-^ macrophages. Although E3 ligases are often highly substrate specific [30], our data suggest that the GID/CTLH complex has exceptional functional versatility in Mtb-infected macrophages, as its loss broadly disrupts global protein homeostasis.

While individual GID/CTLH complex members have been implicated in various immune processes [14], the broad impact of the complex on macrophage immunometabolism had largely gone unreported until the recent Mtb infection studies [25]. In agreement with the observations that Mtb-infected GID/CTLH macrophages exhibit a strongly anti-inflammatory phenotype [25], immune-related pathways were significantly enriched among the decreased proteins in Mtb-infected GID8^-/-^ and MAEA^-/-^ BMDM^h^s (Fig. 3B, S2 Data). Enriched categories included innate immune response, antiviral innate immune response, pattern recognition receptor signaling (including cytoplasmic PRR signaling), canonical NF-κB signaling, and type I interferon-mediated pathways. At the same time, enriched themes among decreased proteins include cellular responses to interferon-β, regulation and production of type I interferons, cytokine-mediated signaling, and defense responses to viral and protozoan stimuli. These results indicate attenuation of innate immune sensing and downstream inflammatory and antiviral effector programs. We also observed a concomitant enrichment of pathways associated with killing and disruption of cells of another organism which further supports reduced cytotoxic and antimicrobial activity in Mtb-infected GID8^-/-^ and MAEA^-/-^ macrophages. Given that GID/CTLH-deficient macrophages robustly control the intracellular growth of Mtb, these data suggest that loss of the complex in the context of Mtb infection results in cytokine-independent control of bacterial growth, in a manner that could define a metabolically enforced paradigm of bacterial control that does not require immune activation [31].

Several metabolic and translational processes were also enriched among decreased proteins, indicating that metabolic reprogramming in these macrophages does not bifurcate cleanly between anabolic or catabolic programs but instead reflects pathway-specific changes. Core glycolytic pathways, including glucose catabolism to pyruvate and NADH/NAD regeneration, were enriched among decreased proteins, suggesting reduced glycolytic flux and altered cytosolic redox metabolism. Although this contrasts with our previous metabolic flux data in Mtb-infected GID/CTLH knockout macrophages [25], we believe this is mainly due to the increased resilience of the mutant macrophages, which may display enhanced functional glycolysis due to improved control of infection, rather than an intrinsic increase in glycolytic flux. Enrichment in nucleotide and nucleoside monophosphate metabolic and salvage pathways among the decreased proteins, including deoxyribonucleotide and ribonucleoside monophosphate metabolism, point to altered nucleotide turnover and biosynthetic capacity. Furthermore, translational control pathways, such as cytoplasmic translation initiation and elongation, rescue of stalled ribosomes, and ribosome disassembly, were enriched among decreased proteins, indicating global dampening of protein synthesis and ribosome dynamics. These changes suggest that, despite enhanced mitochondrial oxidative metabolism, cytosolic anabolic capacity and biosynthetic readiness may be constrained due to increased mitochondrial redirection of metabolic substrates.

Overall, the differential proteomic landscape of Mtb-infected GID8^-/-^ and MAEA^-/-^ BMDM^h^s reveals a coordinated rewiring of cellular metabolism and immune signaling, characterized by enhanced mitochondrial oxidative metabolism and substrate catabolism alongside broad attenuation of innate immune pathways. Although the contribution of the GID/CTLH complex to metabolic control in mammalian cells has remained a point of contention [13, 14, 19], our data demonstrate that, in macrophages, loss of GID/CTLH activity results in extensive metabolic reprogramming, consistent with a central regulatory role of the complex in coordinating bioenergetic states and cellular metabolic functions during infection. The data implicate the GID/CTLH complex as a key integrator of metabolic and innate immune programs in macrophages, enforcing a bioenergetically constrained and immunologically primed state that shapes host-pathogen interactions and the intracellular survival of Mtb under physiological conditions.

### Quantitative diGly proteomics identifies candidate ubiquitylation substrates in Mtb-infected macrophages lacking GID8 or MAEA

To directly identify candidate substrates ubiquitylated by the GID/CTLH complex during Mtb infection, we performed diGly ubiquitin-remnant profiling [32, 33] in Mtb-infected scramble control, GID8^-/-^ and MAEA^-/-^ BMDM^h^s in parallel with global LFQ proteomics (Fig 1A). This approach enabled site-resolved quantification of lysine ubiquitylation events and allowed us to assess how disruption of either the structural scaffold (GID8) or the catalytic core (MAEA) of the complex reshapes the macrophage ubiquitylome during infection. Across all the conditions, we identified 23,841 diGly-modified lysine sites corresponding to 4,268 proteins (Fig 4A, S3 Data), indicating extensive ubiquitylation of host proteins in Mtb-infected macrophages. This dataset, together with a recently published study [34], provides an extremely detailed atlas of ubiquitin modification in Mtb-infected macrophages.

**Fig 4.**
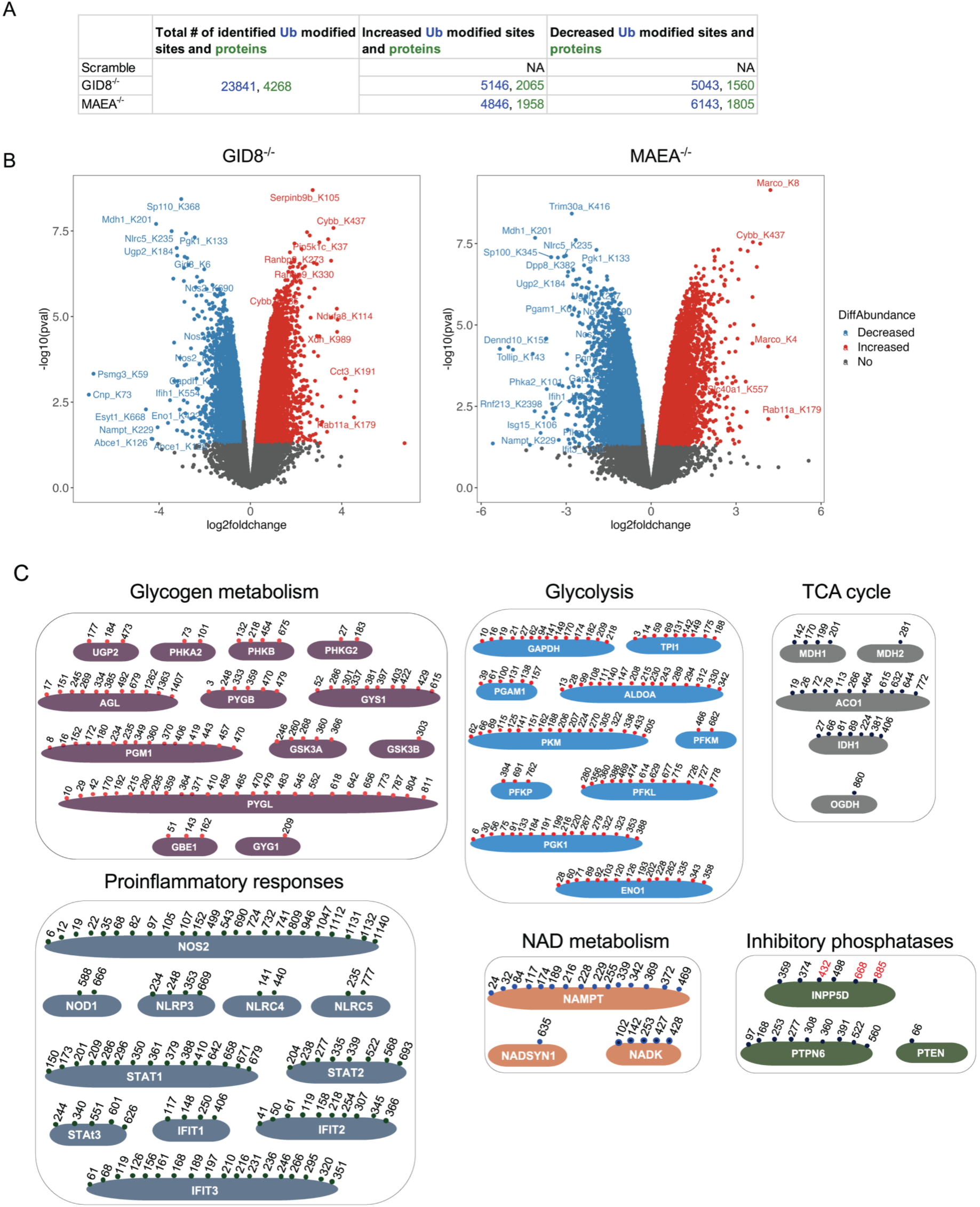
Quantitative diGly proteomics to identify potential ubiquitylation substrates in GID8^-/-^ and MAEA^-/-^ Mtb-infected BMDM^h^s. **A**. All identified diGly-modified peptides and their corresponding proteins detected in the experiment. DiGly-modified peptides and associated proteins showing significant changes in ubiquitylation upon knockout in Mtb-infected GID8^-/-^ and MAEA^-/-^ macrophages are indicated (Data S3). Significant increase or decrease in ubiquitin modified sites abundance was defined as *p* ≤ 0.05 with log₂ fold change ≥ 0.3 (Up) or ≤ −0.3 (Down). **B.** Volcano plots showing changes in diGly-modified peptides in Mtb-infected macrophages upon knockout of GID8 and MAEA. Red and blue dots represent diGly-modified peptides that are at least 1.2-fold increased or decreased in abundance, respectively (log2 ≥ 0.3 and log2 ≤ −0.3, respectively) and have a *p* value of ≤0.05. **C.** Selected proteins across indicated pathways showing commonly decreased diGly modifications (candidate substrates of the GID/CTLH complex) in Mtb-infected GID8^-/-^ and MAEA^-/-^ BMDM^h^s. Numbers indicate modified lysine positions that are shared between the two mutants (black) or unique to MAEA^-/-^ BMDM^h^s (red).

Consistent with the broad proteomic remodeling observed upon GID/CTLH disruption (Fig 1-3), loss of either GID8 or MAEA induced widespread changes in protein ubiquitylation. In GID8^-/-^ BMDM^h^s, 5,146 diGly-modified sites on 2,065 proteins were significantly increased in abundance, whereas 5,043 sites on 1,560 proteins were significantly decreased. Similarly, MAEA^-/-^ BMDM^h^s exhibited 4,846 increased diGly-modified sites on 1,958 proteins and 6,143 decreased sites on 1,805 proteins relative to scramble controls (Fig 4A). The magnitude of these changes underscores a central role for the GID/CTLH complex in shaping the global ubiquitylation landscape of macrophages during Mtb infection. In a given sample, the abundance of a ubiquitylated peptide can change either due to altered stoichiometry of ubiquitylation at a specific site or due to changes in the abundance of the parent protein. However, proteasome inhibition or general proteotoxic stress typically leads to pronounced ubiquitylation across the proteome, but with low overall stoichiometry that does not usually correlate with measurable changes in total protein abundance [34–36]. We therefore chose not to normalize or correlate diGly peptide changes to total protein abundance, as such a correction would obscure biologically meaningful, GID/CTLH-dependent ubiquitylation events.

Volcano plot analyses revealed highly concordant patterns of ubiquitylation changes in GID8^-/-^ and MAEA^-/-^ BMDM^h^s (Fig 4B). This concordance was further supported by extensive overlap of both increased and decreased diGly-modified sites between GID8^-/-^ and MAEA^-/-^ BMDM^h^s (Fig S2A), with ∼59% of increased and ∼54% of decreased ubiquitylation sites shared between the two knockout conditions. Unsupervised hierarchical clustering of differentially abundant diGly-modified peptides demonstrated near-identical global remodeling of the ubiquitylome in GID8^-/-^ and MAEA^-/-^ BMDM^h^s relative to scramble controls (Fig S2B), supporting disruption of a shared functional complex rather than subunit-specific effects. To gain mechanistic insight into pathways directly controlled by GID/CTLH-mediated ubiquitylation, we focused on diGly-modified sites that were consistently and commonly decreased in both GID8^-/-^ and MAEA^-/-^BMDM^h^s, reasoning that these likely represent candidate substrates of the complex (Fig 4B, Fig S2A-B, S3 Data). Strikingly, proteins involved in core metabolic pathways were highly enriched among these putative substrates. Multiple enzymes involved in glycogen metabolism, glycolysis, and the TCA cycle harbored shared and in some instances >20 lysine residues with reduced ubiquitylation in both knockout conditions (Fig 4B-C, S3 Data). These data indicate that the GID/CTLH complex directly targets a broad array of metabolic enzymes for ubiquitylation during Mtb infection, which is in agreement with the extensive metabolic rewiring observed at the proteome level (Fig 3). In addition to metabolic enzymes, several innate immune and inflammatory signaling targets exhibited reduced ubiquitylation in GID/CTLH-deficient macrophages, including components of inflammasome and pattern recognition pathways (NOD1, NLRP3, NLRC4, NLRC5), inducible nitric oxide synthase (NOS2), STAT family transcription factors (STAT1, STAT2, STAT3), and interferon-stimulated gene products (IFIT1-3) (Fig 4C, S3 Data). We also identified proteins involved in NAD metabolism (NAMPT, NADK, NADSYN1) as candidate substrates of the GID/CTLH complex, linking ubiquitin-dependent signaling to control of cellular redox balance and metabolic cofactor availability, which corroborates recent findings implicating the complex in the ubiquitylation of the NAD metabolic enzyme NMNAT1 [16]. Additionally, the inhibitory phosphatases INPP5D (SHIP1), PTPN6 (SHP-1), and PTEN also emerged as prominent candidate substrates (Fig 4C, S3 Data), highlighting inhibitors of the PI3K-AKT signaling pathway as potential GID/CTLH-targeted proteins during Mtb infection.

In agreement with these substrate-level findings, pathway enrichment analysis of proteins harboring commonly decreased diGly-modified sites between the two mutants revealed marked enrichment of metabolic and immune pathways (Fig S3A, S4 Data). Interestingly, among these, pathways involved in vesicle-mediated transport, neuronal development, cell motility, endosomal trafficking, actin cytoskeleton organization, phosphoinositide metabolism, and small GTPase signaling were also enriched. These data extend the functional scope of GID/CTLH-dependent ubiquitylation in Mtb-infected macrophages, highlighting its broad involvement in additional biological pathways, as previously reported in other mammalian cells [13, 14, 20]. Conversely, several diGly-modified peptides and pathways among proteins harboring commonly increased diGly sites were elevated upon GID/CTLH disruption (Fig 4A-B; Fig S2; Fig S3B; S3-S4 Data), likely reflecting secondary remodeling of ubiquitin-dependent signaling networks and compensatory ubiquitylation by other E3 ligases in response to altered proteostasis.

### PTEN and INPP5D are stabilized in GID8^-/-^ and MAEA^-/-^ macrophages in a proteasome-dependent manner

Our diGly ubiquitylome analysis identified numerous candidate substrates whose ubiquitylation was consistently reduced in both GID8^-/-^ and MAEA^⁻/⁻^ BMDM^h^s during Mtb infection (Fig 4B-C; S3 Data), spanning proteins involved in innate immune signaling, NAD metabolism, and central carbon metabolism. To functionally validate whether these changes in ubiquitylation reflect altered protein turnover, we performed cycloheximide (CHX) chase assays on a representative set of candidate substrates, including NLRP3 (immune signaling), NAMPT (NAD biosynthesis), and the metabolic enzymes MDH1, MDH2, and UGP2. In scramble control BMDM^h^s, the inflammasome protein NLRP3 exhibited measurable turnover over the 24-hour CHX chase period, whereas in GID8^⁻/⁻^ and MAEA^⁻/⁻^ BMDM^h^s there was evidence of delayed degradation, resulting in significantly higher residual protein levels at 12 and 24 hours following CHX treatment (Fig S4A). In contrast, we did not observe differences in degradation kinetics for NAMPT, UGP2, MDH1, or MDH2 (Fig S4A-C). Notably, MDH2 was among the earliest substrates identified for the GID complex in *Saccharomyces cerevisiae* and is now a well-characterized target of GID-mediated degradation [11, 37]. These data suggest that loss of the GID/CTLH complex selectively stabilizes a subset of substrates in mammalian macrophages, including immune signaling proteins such as NLRP3. For the other diGly-identified candidates tested, the absence of detectable changes in protein abundance over the CHX chase may reflect longer intrinsic half-lives under these conditions and/or ubiquitylation events that do not target proteins for proteasomal degradation.

We next focused on three inhibitory phosphatases, PTPN6 (SHP-1), INPP5D (SHIP1), and PTEN, which emerged as high-confidence candidate substrates in the diGly dataset (Fig 4C; S3 Data) and were also identified in our previous CRISPR-Cas9 knockout screen [25] as potential candidates that could restrict intracellular Mtb growth in macrophages. In the CHX chase assay, we did not observe differences in PTPN6 degradation kinetics over the 24-hour period (Fig S4D). INPP5D, however, was rapidly degraded in scramble control macrophages, with protein levels declining sharply within 12 hours (Fig 5A), but was significantly stabilized in both GID8^⁻/⁻^ and MAEA^⁻/⁻^ BMDM^h^s, with ∼50-100% of the initial protein abundance remaining at 12-24 hours (Fig 5A-B; Fig S5A-C). Steady-state INPP5D protein levels were also elevated in both knockout genotypes relative to scramble controls (Fig 5A, Fig S5A-B) suggesting that reduced ubiquitylation of INPPD5 reduces its targeting to the proteasome and increases its levels and stability in GID/CTLH-deficient macrophages. To determine whether INPP5D degradation is proteasome-dependent and whether this process requires the GID/CTLH complex, scramble control and MAEA^⁻/⁻^ BMDM^h^s were treated with CHX or the proteasome inhibitor MG132 in separate conditions for 18 hours. In scramble control cells, CHX treatment led to a marked reduction in INPP5D protein levels (Fig 5B, Fig S5C), in line with our prior observations (Fig 5A; Fig S5A-B), whereas MG132 treatment alone significantly increased INPP5D abundance, consistent with proteasome-mediated turnover (Fig 5B, Fig S5C). In contrast, INPP5D levels in MAEA^⁻/⁻^ BMDM^h^s remained relatively stable following CHX treatment, and MG132 treatment produced no additional accumulation of INPP5D (Fig 5B, Fig S5C), indicating that loss of MAEA attenuates proteasome-dependent degradation of INPP5D.

**Fig 5.**
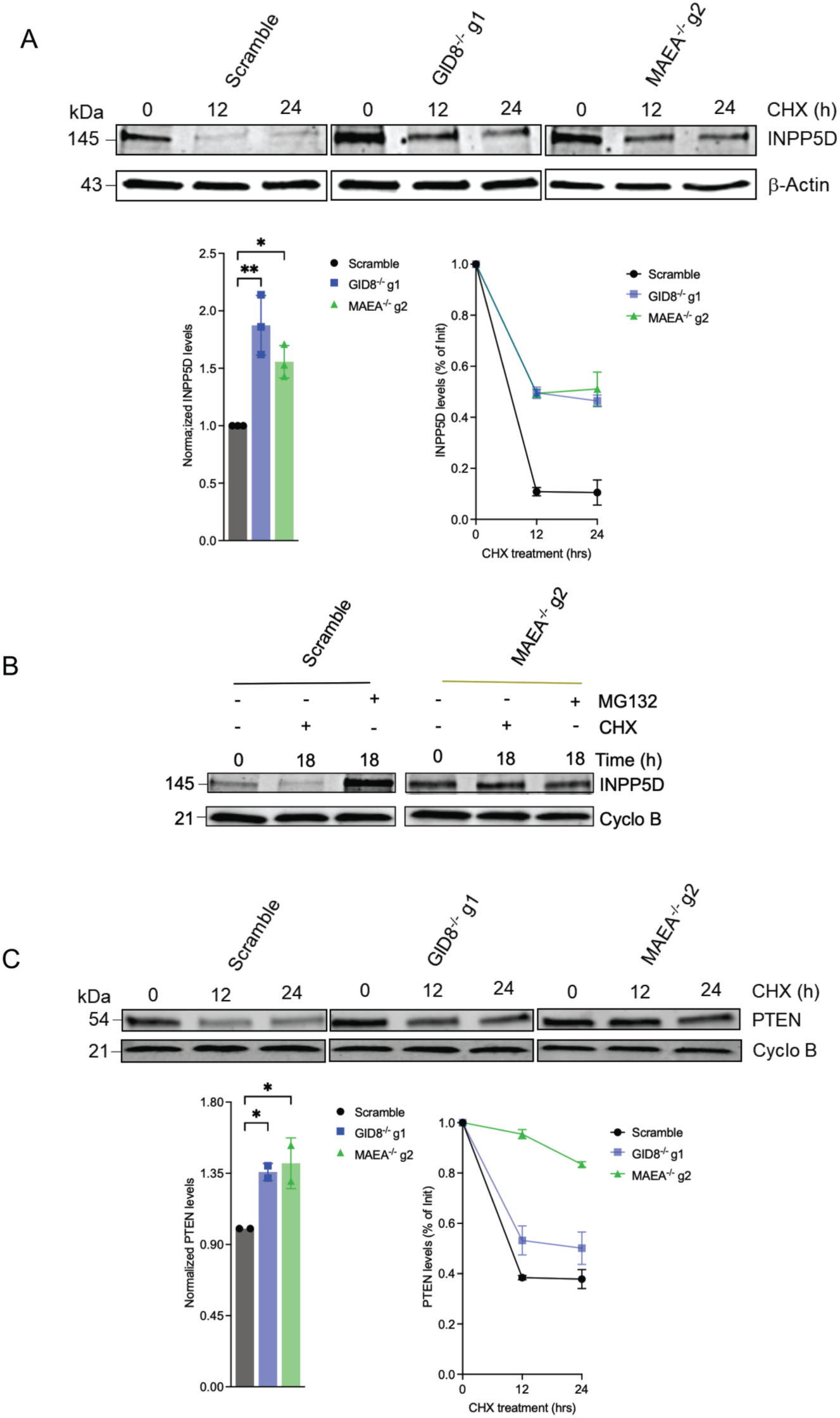
Proteasome dependent stabilization of PTEN and INPP5D protein levels in GID8^-/-^ and MAEA^-/-^BMDM^h^s. **A.** Cycloheximide (CHX) chase assay performed to monitor the degradation of INPP5D in scramble control, GID8^-/-^, and MAEA^-/-^ BMDM^h^s over a 24-hour period. Protein levels were analyzed by immunoblotting. Normalized protein levels and the percentage of protein remaining in reference to initial levels at the indicated time points are plotted for each protein. **B**. Immunoblot analysis shows that INPP5D degradation is proteasome-dependent, as CHX treatment leads to protein degradation, while the proteasome inhibitor MG132 results in a significant increase in basal INPP5D levels in scramble control BMDM^h^s after 18 hours of treatment. In contrast, INPP5D exhibits increased stabilization in CHX-treated MAEA^-/-^ BMDM^h^s, and MG132 does not further increase INPP5D protein levels in this mutant. **C.** CHX chase assay to monitor the degradation of INPP5D in scramble control, GID8^-/-^, and MAEA^-/-^ BMDM^h^s over a 24-hour period, performed as in (A). Quantification of relative protein expression was performed for replicate blots shown in (A) and (C) and in Fig. S5A-S5D, normalized to either β-actin or Cyclophilin B. Data represent n = 3 biological replicates for INPP5D and n = 2 biological replicates for PTEN. *P < 0.05; **P < 0.01, two-way ANOVA alongside Dunnett’s multiple comparison test. Data are presented as mean values ± SD.

PTEN exhibited similar stabilization kinetics in the CHX chase assay, with significantly delayed degradation and increased steady-state protein levels in GID8^⁻/⁻^ and MAEA^⁻/⁻^ BMDM^h^s compared to scramble controls (Fig 5C; Fig S5D). Notably, ∼40% of PTEN protein remained after 24 hours of CHX treatment, in contrast to INPP5D (<15%), suggesting that PTEN could be ubiquitylated by additional ubiquitin ligases beyond the GID/CTLH complex. Consistent with this possibility, several E3 ligases have indeed been reported to ubiquitylate PTEN in mammalian cells, including NEDD4-1, WWP2, CHIP, XIAP, RFP, HRD1, and the CUL4B–DCAF13 complex [38].

### Genetic or pharmacological inhibition of INPP5D and PTEN impairs Mtb replication in macrophages

We identified PTEN and INPP5D as potential substrates of the GID/CTLH complex and confirmed their increased stability and proteasome-dependent degradation in BMDM^h^s lacking individual members of the complex (Fig 4-5). Both proteins are inhibitors of PI3K-Akt signaling and modulate pro-inflammatory responses in macrophages [39, 40]. Because GID/CTLH-deficient BMDM^h^s display a strongly anti-inflammatory phenotype [25]; it is unlikely that increased stabilization of PTEN and INPP5D contributes to enhanced Mtb growth control in these cells. However, if these proteins are routinely degraded by the proteasome under physiological conditions in macrophages, potentially to relieve inhibition of pro-inflammatory signaling, we reasoned that their knockout in wild-type macrophages should enhance PI3K-Akt signaling, promote pro-inflammatory cytokine production, and restrict intracellular Mtb growth. We therefore investigated whether PTEN or INPP5D modulates intracellular Mtb replication, by generating independent CRISPR-Cas9 knockout Hoxb8 macrophage lines for each target (S5 Data). Immunoblot analysis confirmed efficient depletion of PTEN and INPP5D proteins in the respective knockout macrophages relative to scramble controls (Fig 6A).

**Fig 6.**
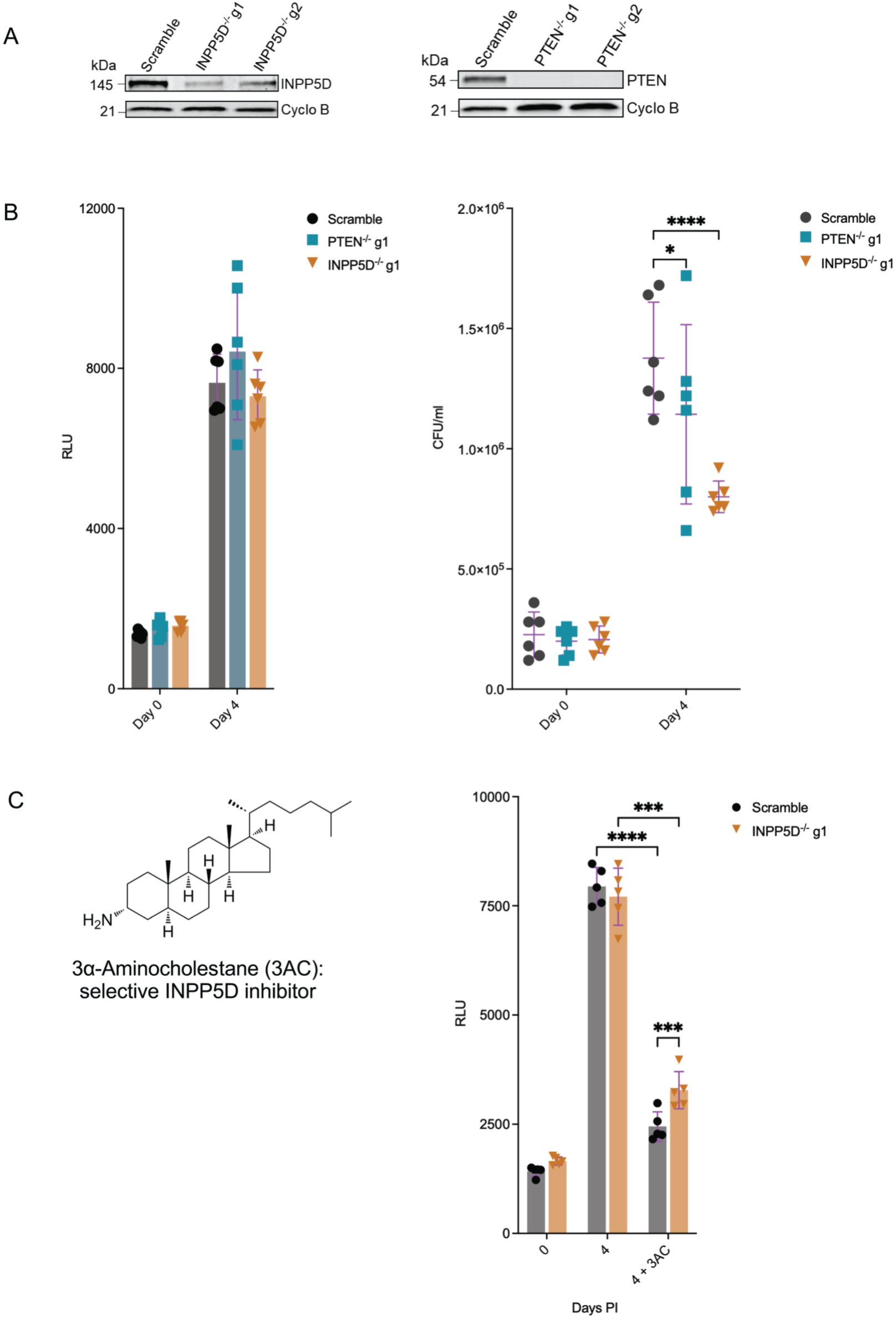
INPP5D and PTEN inhibition, either chemical or genetic, limits Mtb growth in BMDM^h^s. **A**. Immunoblot confirmation of target protein depletion (INPP5D and PTEN) in generated knockout BMDM^h^s derived from Hoxb8 parental mutant lines described in Data S5. **B.** Mtb growth kinetics in INPP5D^-/-^ and PTEN^-/-^ BMDM^h^s compared with scramble controls as measured by luciferase activity or colony-forming units (CFUs). For the luciferase-based growth assay, scramble control, INPP5D^-/-^, or PTEN^-/-^ BMDM^h^s were infected with the Mtb Erdman-Lux strain at MOI 0.5, and luciferase measurements were taken on the indicated days (n = 6 biological replicates). For the CFU-based growth assay, mutant BMDM^h^s were infected with Mtb Erdman strain at MOI 0.4. CFUs were enumerated on day 0 (3 hours post infection) and on Day 4 to assess intracellular Mtb replication rates (n = 6 biological replicates). **C.** Activity of the INPP5D inhibitor 3α-aminocholestane (3AC) against Mtb in scramble control and INPP5D^-/-^ BMDM^h^s. The chemical structure of 3AC is shown. BMDM^h^s were infected with the Mtb Erdman-Lux strain at MOI 0.5, and drug treatment was initiated 3-4 hours post-infection. Luciferase measurements were carried out on day 0 and day 4 (n = 5 biological replicates). \**P* < 0.05; \*\*\**P* < 0.001; \*\*\*\**P* < 0.0001, two-way ANOVA alongside Dunnett’s multiple comparison test. Data are presented as mean values ± SD.

We next quantified intracellular Mtb growth in PTEN^-/-^ and INPP5D^-/-^ BMDM^h^s using both luminescence-based bacterial growth assays following infection with Mtb-Lux and standard CFU enumeration following infection with wild-type Mtb Erdman. By the luciferase-based readout, neither PTEN^-/-^ nor INPP5D^-/-^BMDM^h^s exhibited a detectable difference in bacterial growth compared with scramble controls at day 4 post infection (Fig 6B), in line with observations that luciferase readouts lack the dynamic range to resolve more modest bacterial control phenotypes as we previously observed with lipid knockout mutant BMDM^h^s [41] and preferentially detects strong restriction phenotypes such as those observed upon disruption of the GID/CTLH complex itself [25]. In contrast, CFU enumeration revealed a reduction in intracellular bacterial growth in both INPP5D^-/-^ and PTEN^-/-^ BMDM^h^s at day 4 post infection relative to scramble controls (Fig 6B), with a more pronounced and consistent effect observed in INPP5D-deficient BMDM^h^s and a more modest effect in PTEN-deficient macrophages. Importantly, bacterial uptake at day 0 was comparable between genotypes, indicating that differences in bacterial burden at later time points reflected intracellular growth control rather than altered phagocytic uptake.

To further validate the functional role of INPP5D activity, we pharmacologically inhibited INPP5D using the selective small-molecule inhibitor 3α-aminocholestane (3AC) [42] in scramble control and INPP5D^-/-^BMDM^h^s infected with Mtb-Lux. While we found that the inhibitor restricted Mtb growth under both macrophage conditions (Fig 6C), we also observed that the compound was active against Mtb in broth culture (Fig S6). Given the inhibitor’s structure (Fig 6C), this is perhaps not surprising due to Mtb’s known susceptibility to inhibitors of bacterial cholesterol metabolism [43, 44]. We observed a partial loss of 3AC inhibitory activity against Mtb in INPP5D^-/-^ BMDM^h^s (Fig 6C), which supportively confirms the inhibitor specificity. Nonetheless, the genetic knockouts confirm the roles of PTEN and INPP5D as functionally relevant host factors that individually limit macrophage antimicrobial capacity against Mtb, albeit with more modest effects than loss of the GID/CTLH complex itself.

### Loss of INPP5D activity increases Akt phosphorylation and promotes a pro-inflammatory response in Mtb-infected macrophages

Next, we assessed the functional consequences of INPP5D loss or inhibition on downstream signaling pathways in infected BMDM^h^s, as INPP5D displayed a stronger antibacterial phenotype than PTEN and is predominantly expressed in immune cells [45]. First, we examined Akt phosphorylation in both uninfected and Mtb-infected BMDM^h^s treated or not treated with 3AC, in line with the established role of INPP5D as an inhibitor of PI3K-Akt signaling in macrophages [39]. INPP5D^-/-^ BMDM^h^s displayed increased levels of phosphorylated Akt (p-Akt) in the uninfected state compared with scramble controls, while total Akt levels remained unchanged (Fig 7A, Fig S7A). Interestingly, Mtb infection also increased p-Akt levels in scramble control BMDM^h^s, but the level of phosphorylation was even more pronounced in Mtb-infected INPP5D^-/-^BMDM^h^s. Pharmacologic inhibition of INPP5D using 3AC similarly enhanced Akt phosphorylation in infected INPP5D^-/-^ BMDM^h^s but was relatively blunted in scramble control macrophages. These results indicate that INPP5D knockout enhances PI3K-Akt pathway activation during Mtb infection and that the 3AC inhibitor shows some evidence of INPP5D target engagement in infected BMDM^h^s, but not at the same level as the genetic knockout.

**Fig 7.**
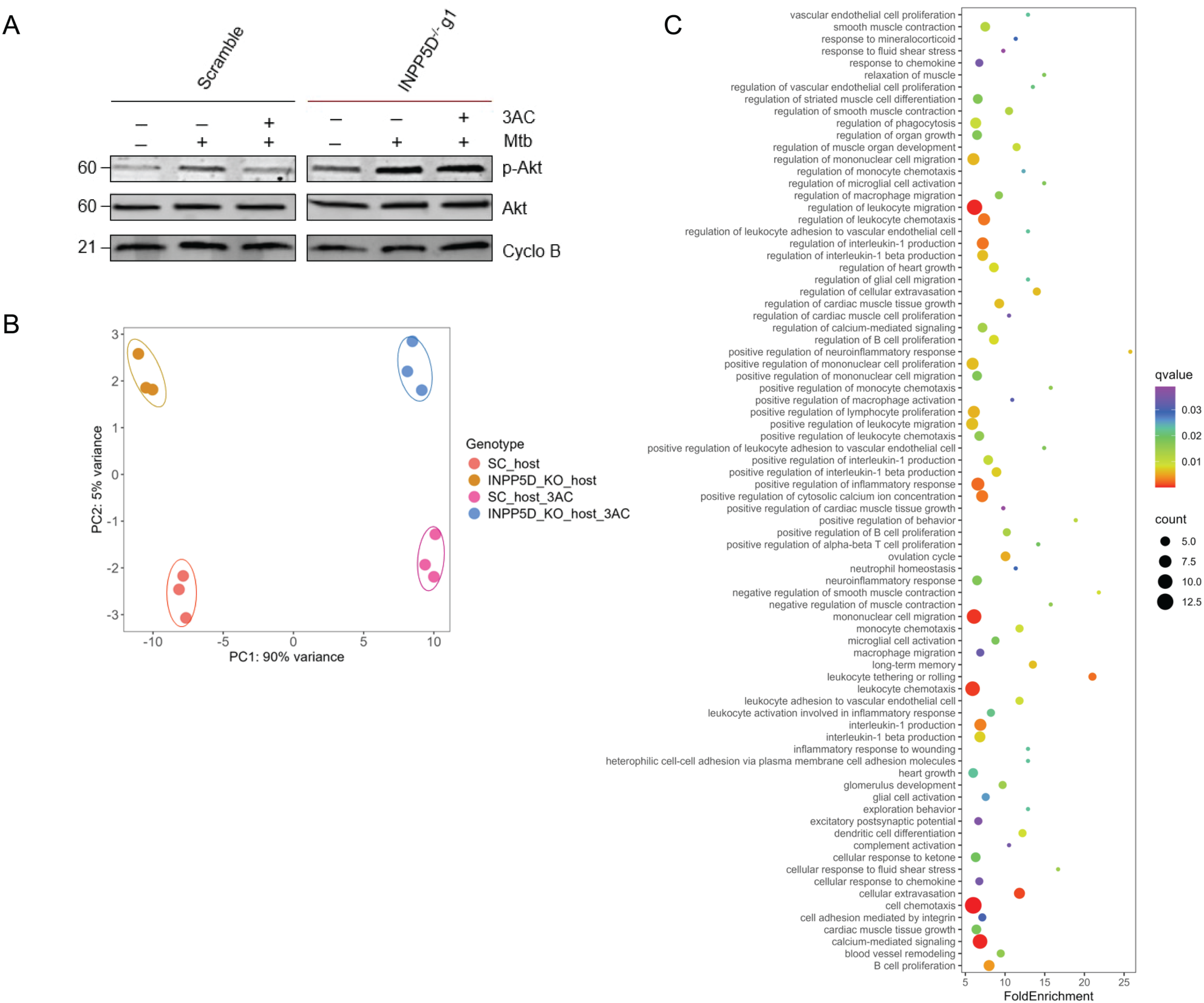
Impairment of INPP5D activity increases Akt phosphorylation and promotes a pro-inflammatory response in Mtb-infected BMDM^h^s. **A**. Immunoblot analysis of total Akt and phosphorylated Akt (p-Akt) in scramble control and INPP5D^-/-^ BMDM^h^s. Cells were left uninfected or infected with Mtb Erdman at an MOI of 3 for 24 hours and treated or untreated with 3AC beginning 3 hours post-infection, as indicated. **B.** Principal component analysis (PCA) of host transcriptional responses in scramble control or INPP5D^-/-^BMDM^h^s, untreated or treated with 3AC, following infection with the Mtb smyc’::mCherry strain at MOI 0.5 for 4 days. **C**. Top enriched biological pathways among upregulated genes in Mtb-infected INPP5D^-/-^macrophages relative to scramble controls.

We performed transcriptomic profiling of Mtb-infected scramble control and INPP5D^-/-^ BMDM^h^s treated or untreated with 3AC to identify INPP5D-dependent pathways of bacterial control in infected macrophages. PCA analysis revealed clear segregation of Mtb-infected BMDM^h^s by both genotype and chemical INPP5D inhibition (Fig 7B), demonstrating that genetic and pharmacologic disruption of INPP5D elicit robust and distinct host transcriptional responses to infection. Pathway enrichment analysis of genes upregulated in infected INPP5D^-/-^ BMDM^h^s (S6 Data) showed strong enriched of inflammatory and immune response programs, including leukocyte chemotaxis and migration, leukocyte adhesion and extravasation, interleukin-1 production, cytokine-mediated signaling, and regulation of inflammatory responses (Fig 7C, S7 Data). These findings confirm that INPP5D functions as an inhibitor of pro-inflammatory gene expression in Mtb-infected BMDM^h^s, and that its deletion enhances these responses to restrict intracellular bacterial growth. The results are also in agreement with recent reports implicating loss of INPP5D with enhanced inflammasome activation in microglia [46]. Pathways enriched among downregulated genes (S6 Data) in Mtb-infected INPP5D^−/−^ BMDM^h^s were primarily related to general kinase and cellular signaling (Fig S7B, S7 Data) and did not reveal additional biologically informative programs relevant to macrophage antimicrobial programs. Mtb-infected scramble control and INPP5D^−/−^ BMDM^h^s treated with 3AC displayed comparable patterns of pathway enrichment, characterized by upregulated cholesterol metabolism, acyl-CoA metabolic processes, and steroid biosynthetic processes, alongside downregulated pro-inflammatory responses, including inflammasome activation and cytokine production (Fig S8-S9, S6-S7 Data). This is consistent with the strong lipid-modulatory activity of 3AC, a cholesterol derivative (Fig 6C), as well as its direct bactericidal activity (Fig S6).

## Discussion

In this study, we demonstrate that the GID/CTLH E3 ubiquitin ligase complex is a central regulator of macrophage metabolism and antimicrobial immunity during Mtb infection. By integrating global proteomics with ubiquitin-remnant profiling, we reveal that loss of GID/CTLH activity induces extensive remodeling of both protein abundance and ubiquitylation across metabolic and immune signaling pathways. Our data implicate the GID/CTLH complex as a systems-level coordinator of macrophage cellular metabolic state, linking ubiquitin-dependent signaling to bioenergetic programming and intracellular bacterial control.

A striking finding of our study is the dominant metabolic signature induced by loss of GID/CTLH activity during Mtb infection of macrophages. GID8^-/-^ and MAEA^-/-^ macrophages displayed broad upregulation of mitochondrial oxidative phosphorylation, fatty acid β-oxidation, and TCA cycle pathways, along with increased abundance of mitochondrial gene expression programs and protein import machinery across the global proteome. These changes indicate a shift toward enhanced mitochondrial respiratory capacity and substrate oxidation. This metabolic rewiring agrees with our prior functional observations that GID/CTLH-deficient macrophages maintain metabolic resilience during Mtb infection and impose oxidative stress on intracellular bacteria [25]. For many years, it has been recognized that macrophages possess remarkable metabolic plasticity, and that their metabolic programs often mirror the disease environments in which they reside [47, 48]. Historically, these contrasting metabolic phenotypes were largely attributed to external environmental cues, such as tissue-derived signals and the surrounding cytokine landscape. This perspective was influenced by the long-standing assumption that all macrophages originate from circulating blood monocytes and exist in a basal “M0” state that can be polarized into M1 or M2 phenotypes in response to specific cytokines [47–49]. However, more recent insights into macrophage heterogeneity, encompassing both developmental origin and pre-programmed metabolic states, have revealed a far more complex landscape than previously appreciated [3, 50, 51]. During intracellular infection with Mtb in the lung, for example, resident alveolar macrophages respond differently from recruited interstitial macrophages, even when exposed to identical cytokine environments. These divergent responses arise from differences in ontogeny and intrinsic metabolic wiring, which together shape each macrophage population’s capacity to support or restrict bacterial growth [50, 52]. This complexity raises an important question: is there a key regulator that coordinates macrophage metabolism across disparate developmental, environmental, and tissue conditions? Our data identify the GID/CTLH complex as a central regulator of cellular metabolism in Mtb-infected macrophages, and a potential candidate for such a role. The dynamic nature of ubiquitylation itself may confer the flexibility required to fine-tune metabolic pathways, ensuring that only the appropriate metabolic programs are engaged, whether during adaptive re-programming or within pre-established metabolic states, to support context-specific cellular responses. The findings support a model in which GID/CTLH activity constrains mitochondrial bioenergetic capacity under basal and infection-induced conditions, thereby limiting macrophage metabolic fitness during infection. In this context, loss of the GID/CTLH complex in knockout conditions appears to release this bioenergetic brake, enabling macrophages to sustain enhanced oxidative metabolism and better tolerate infection-induced stress.

Our diGly ubiquitin-remnant profiling revealed that the GID/CTLH complex directly targets a remarkably wide spectrum of substrates, including enzymes spanning glycolysis, glycogen metabolism, the TCA cycle, and NAD metabolism. This extensive targeting of central carbon metabolism enzymes suggests that the GID/CTLH complex may function as a master regulator of metabolism in macrophages, shaping pathway fluxes and substrate availability through ubiquitin-dependent modulation of enzyme turnover and/or non-degradative ubiquitin signaling. Although the role of the mammalian GID/CTLH complex in metabolic control remains a subject of discussion [13, 14, 19], our data provide direct *in situ* evidence in primary macrophages that the complex exerts broad and coordinated control over metabolic networks during Mtb infection. Notably, not all diGly-identified candidate substrates exhibited altered proteasomal degradation kinetics, suggesting that GID/CTLH-dependent ubiquitylation likely serves both degradative and non-degradative functions, consistent with emerging evidence of increased versatility in ubiquitin-mediated signaling [9, 30].

It is significant that enhanced control of intracellular Mtb in GID/CTLH-deficient macrophages occurs despite broad suppression of canonical inflammatory and cytokine-driven antimicrobial programs. This phenotype challenges the prevailing paradigm that macrophage-mediated control of Mtb is dependent solely on inflammatory activation [31] and highlights an alternative mode of bacterial restriction with the potential to be cytokine independent. We believe that metabolic fitness and improved mitochondrial function constitute a dominant layer of antimicrobial control in this context. Enhanced oxidative metabolism, redox capacity, and substrate catabolism may promote intrinsic host cell resilience and create a metabolically restrictive intracellular environment that limits Mtb replication. This closely aligns with emerging concepts of nutritional immunity and metabolically enforced host restriction of intracellular pathogens [3, 41, 50, 51].

Among the GID/CTLH-ubiquitylated substrates, we identified the inhibitory phosphatases PTEN and INPP5D as proteasome-degraded targets whose stability is increased in GID8^-/-^ and MAEA^-/-^ macrophages. Both proteins are inhibitors of PI3K-Akt signaling, a pathway that has been implicated in macrophage survival, polarization and inflammatory activation [39, 40], and were also lead host candidates identified in our CRISPR-Cas9 knockout screen that uncovered the GID/CTLH complex [25]. Genetic loss of PTEN or INPP5D in wild-type macrophages modestly restricted intracellular Mtb growth, with INPP5D exerting the stronger effect, and INPP5D deficiency enhanced Akt phosphorylation and pro-inflammatory transcriptional responses during infection. These findings suggest that basal GID/CTLH-dependent turnover of inhibitory phosphatases allows for a dynamic modulation of PI3K-Akt signaling and inflammatory responsiveness in macrophages [39, 40, 46]. This agrees with our integrated proteomics data, which showed GID/CTLH candidate substrates in several immune pathways and a broader downregulation of pro-inflammatory processes in infected GID/CTLH-deficient macrophages. However, the anti-inflammatory phenotype of GID/CTLH-deficient macrophages indicates that stabilization of PTEN and INPP5D is not the mechanism driving enhanced Mtb control in these cells. Instead, these substrates likely contribute to fine-tuning of inflammatory signaling thresholds under physiological conditions, whereas the robust antibacterial phenotype observed upon GID/CTLH disruption reflects the combined effect of widespread metabolic and proteostatic perturbation.

Several considerations are relevant to the interpretation of our findings. First, diGly profiling identifies candidate substrates but does not distinguish between direct and indirect targets of ubiquitylation, nor does it resolve the specific ubiquitin chain topologies involved [33, 35, 53, 54]. Second, many ubiquitylation events likely mediate non-degradative signaling functions [55, 56], which may require modification-specific antibodies to validate, and were not systematically assessed. Third, our experiments were performed in *ex vivo* differentiated Hoxb8 derived bone marrow primary macrophages; whether similar modulatory functions operate in primary macrophage populations *in vivo* during Mtb infection remains to be established. These limitations do not undermine the relevance of our findings; however, they highlight important directions for future work to refine the mechanistic understanding and physiological relevance of GID/CTLH-dependent ubiquitin signaling in broader host-pathogen interactions.

In summary, our findings identify the GID/CTLH E3 ligase complex as a central regulator of ubiquitin-dependent control of macrophage metabolism and antimicrobial immunity during Mtb infection. By constraining mitochondrial bioenergetic capacity and shaping the ubiquitylation landscape of metabolic and immune signaling proteins, the GID/CTLH complex enforces a cellular state that favors Mtb persistence under basal and infection-induced conditions in macrophages. Disruption of this function through knockout of subunit members of the GID/CTLH complex results in a metabolically enforced mode of Mtb control that is largely independent of canonical inflammatory pathways, highlighting ubiquitin-mediated regulation of host metabolism as an underappreciated determinant of antimicrobial responses.

## Materials and methods

### Mtb strains

All CFU and standard infection assays were performed using the parental, PDIM-positive, Mtb Erdman strain (ATCC 35801). Reporter strains in the Erdman background included the fluorescent Mtb Erdman smyc’::mCherry, previously described [57], and the bioluminescent Erdman-Lux strain, generated by transformation with the pMV306G13+Lux plasmid [58]. Mtb cultures were grown to logarithmic phase at 37 °C in Middlebrook 7H9 broth supplemented with 10% OADC enrichment (oleic acid, albumin, dextrose, and catalase; Becton, Dickinson and Company), 0.2% glycerol, and 0.05% tyloxapol (Sigma-Aldrich). Reporter strains were maintained under antibiotic selection as appropriate: the Erdman-Lux strain was cultured with 25 μg/mL kanamycin, and the smyc’::mCherry strain with 50 μg/mL hygromycin.

### Maintenance and culture conditions of mammalian cell lines

Murine Hoxb8 estrogen responsive (ER), Cas9-expressing eGFP⁺ conditionally immortalized myeloid progenitor cells were generated as previously reported [59] and maintained as described extensively elsewhere [25, 41, 60]. Briefly, Hoxb8-ER progenitors were cultured in RPMI medium (Corning®) supplemented with 10% fetal bovine serum (FBS), 2 mM L-glutamine, 1 mM sodium pyruvate, 20 ng/mL recombinant murine GM-CSF (PeproTech), 0.5 μM β-estradiol, 10 mM HEPES, and 1% penicillin/streptomycin. To generate BMDM^h^s, β-estradiol was removed by washing cells twice with 1× PBS. Cells were then transferred to macrophage differentiation medium consisting of DMEM (Corning®) supplemented with 10% FBS, 15% L-cell-conditioned medium, 2 mM L-glutamine, 1 mM sodium pyruvate, and 1% penicillin/streptomycin. Differentiation was carried out at 37 °C for 6-7 days, maintaining cultures at an approximate density of 0.5 × 10⁶ cells/mL throughout the process.

### Generation of mutant CRISPR-Cas9 mutant BMDM^h^s

GID/CTLH-deficient mutant BMDM^h^s used in this study were generated as previously described [25]. All other mutants (PTEN^-/-^ and INPP5D^-/-^) were generated and maintained using lentiviral-delivered CRISPR-Cas9 approaches in Cas9-expressing eGFP⁺ Hoxb8 ERs myeloid progenitors, as described in detail elsewhere [25, 41].

### Macrophage infection assays

BMDM^h^s were infected with Mtb using protocols as previously described [25, 41, 60, 61]. In brief, log-phase cultures were harvested by centrifugation and resuspended in basal uptake buffer composed of 25 mM dextrose, 0.5% bovine serum albumin, 0.1% gelatin, 1 mM CaCl₂, and 0.5 mM MgCl₂ in PBS. To disrupt bacterial aggregates and obtain a single-cell suspension, the preparation was repeatedly passed through a tuberculin syringe (≥20 strokes). Following mechanical dispersion, bacteria were diluted into antibiotic-free macrophage culture medium and added to BMDM^h^s at appropriate MOIs.

### Quantitative proteomics and ubiquitylomics

#### Macrophage infection and protein lysate preparation

Approximately 1 × 10⁸ BMDM^h^s per biological replicate (in triplicate: scramble control, GID8^-/-^, and MAEA-^/-^) were infected with Mtb Erdman at an MOI of 1.5 for 48 hours. After 48 hours, cells were harvested by scraping in ice-cold 1× PBS. Infected cells were sterilized to completely inactivate Mtb by overnight incubation in 90% methanol. The following day, methanol-fixed cells were removed from the BSL3 facility, washed twice with 1× PBS, and resuspended in lysis buffer (8 M urea, 150 mM NaCl, 50 mM TEAB, 1× Halt protease and phosphatase inhibitors). Resuspended cell lysates were sonicated on ice using a large-tip probe at 40% amplitude for three 10-second cycles, with 30-second intervals between cycles (5 seconds on, 5 seconds off). Sonicated lysates were clarified by centrifugation at 14,000 rpm for 30 minutes at 4 °C. Supernatants were transferred to new tubes and protein concentration was measured using the Pierce™ Rapid Gold BCA Protein Assay Kit with a 2 mg/mL albumin standard dilution series. Lysates were stored at −80 °C until shipment to Creative Proteomics (Shirley, New York) where protein analysis was performed. Approximately 5 mg of total protein was used as input for each replicate.

### Sample preparation for proteomics

Cell lysates were subjected to both global LFQ proteomic analysis and diGly profiling. Samples stored at −80 °C were thawed on ice prior to processing. Protease and deubiquitinase inhibitors were added (1 mM PMSF and 1% PR-619; Sigma-Aldrich), followed by vortexing and centrifugation at 4,500 × g for 10 min at 4 °C. Supernatants were collected, and protein concentrations were re-measured using a BCA assay.

### Trypsin digestion

Equal amounts of protein were adjusted to 200 μL in 8 M urea. Samples were reduced with 5 mM dithiothreitol (DTT) at 37 °C for 45 min and alkylated with 11 mM iodoacetamide in the dark at room temperature for 15 min. Proteins were precipitated with four volumes of cold acetone at −20 °C for 2 hours, pelleted by centrifugation, air-dried, and resuspended in 25 mM ammonium bicarbonate. Resuspended protein samples were digested overnight at 37 °C with sequencing-grade trypsin (Promega). Digested peptides were desalted using C18 solid-phase extraction columns (Millipore), quantified using the Pierce™ Quantitative Peptide Assay kit (Thermo Fisher Scientific), concentrated, and vacuum-dried prior to LC-MS/MS analysis. An aliquot of the peptides was used for LFQ proteomics, while the rest were subjected to diGly immunoaffinity enrichment.

### Enrichment of diGly modified peptides

For ubiquitin remnant profiling, digested peptides were subjected to immunoaffinity enrichment of di-glycine (K-ε-GG)-modified peptides using anti-diglycine-lysine antibody-conjugated agarose beads (PTM Biolabs). Peptides were dissolved in enrichment buffer (80% acetonitrile/6% trifluoroacetic acid saturated with glycine) and incubated with pre-washed antibody-conjugated beads on a rotating platform for 1-2 hours at 4°C. After incubation, beads were washed sequentially with 50% acetonitrile/6% trifluoroacetic acid/200 mM NaCl and 30% acetonitrile/0.1% trifluoroacetic acid. Bound peptides were eluted with 10% ammonia solution, vacuum-dried, desalted using C18 ZipTips, and quantified prior to LC-MS/MS analysis.

### Nano LC-MS/MS Analysis

Peptides were separated using a NanoElute UHPLC system (Bruker) equipped with an IonOpticks C18 analytical column (75 μm × 25 cm, 1.6 μm particle size). Approximately 200 ng of peptide was loaded per injection. Mobile phase A consisted of 0.1% formic acid in water, and mobile phase B consisted of 0.1% formic acid in acetonitrile. Peptides were separated at 300 nL/min using a linear gradient: 2-22% B over 45 min, 22-35% B over 5 min, 35-80% B over 5 min, followed by 80% B for 5 min (total runtime: 60 min). Mass spectrometry data was acquired on a timsTOF Pro2 mass spectrometer (Bruker) operating in positive ion mode with the data-dependent acquisition parallel accumulation-serial fragmentation (ddaPASEF) method. The m/z range was 100-1700, and ion mobility (1/K₀) was set from 0.7 to 1.4 Vs/cm². Ion accumulation and release times were 100 ms. Instrument parameters included a capillary voltage of 1,500 V, drying gas flow of 3 L/min, and drying temperature of 180 °C. PASEF settings included 10 MS/MS scans per cycle (cycle time ∼1.17 s), charge range 0-5, dynamic exclusion of 0.4 min, ion target intensity of 10,000, and intensity threshold of 2,500. Collision energy was linearly ramped based on ion mobility (20 eV at 1/K₀ = 0.6 Vs/cm² to 59 eV at 1/K₀ = 1.6 Vs/cm²). Quadrupole isolation widths were set to 2 Th for m/z < 700 and 3 Th for m/z > 800.

### Database Search and quantification

Raw data were processed using FragPipe (v21.1) [62, 63]. Spectra were searched against a combined FASTA database (58,907 sequences) supplemented with decoy and contaminant sequences to control false discovery rates (FDR). Precursor and fragment mass tolerances were set to 20 ppm. For global proteomics, trypsin was specified as the digestion enzyme with up to two missed cleavages permitted. Carbamidomethylation of cysteine was set as a fixed modification, and oxidation of methionine and protein N-terminal acetylation were included as variable modifications. For ubiquitylomics, trypsin and Lys-C were specified as proteases with up to two missed cleavages allowed. Carbamidomethylation (C) was set as a fixed modification, and oxidation (M), protein N-terminal acetylation, and ubiquitylation (diGly modification on K) were included as variable modifications, with a maximum of three variable modifications per peptide. The minimum site localization probability for ubiquitylation sites was set to 0.75. Peptide- and protein-level FDRs were controlled at 1%, and decoy and contaminant hits were excluded from downstream analyses. LFQ was performed using IonQuant with match-between-runs (MBR) and MaxLFQ enabled.

### Differential abundance and ubiquitylation analysis

For LFQ global proteomics, 6,239 proteins matching the mouse protein database were identified. For ubiquitin remnant profiling, 31,749 ubiquitylation sites were detected. After filtering for proteins and peptides identified in all replicates of at least one condition (scramble control, GID8^-/-^ and MAEA^-/-^), a total of 5,087 proteins were retained in the LFQ global proteomics dataset. After similar filtering, 23,841 ubiquitylation sites across 4,268 proteins were retained in the ubiquitylation profiling dataset. Differentially abundant proteins and ubiquitylated peptides were defined as those with absolute fold change (FC) > 1.3 (Up) or < 1.3 (Down) and p < 0.05. Statistical comparison and downstream data analysis was carried out using the proteomic data analysis package DEP in R [64].

### Immunoblot analysis

Cells were harvested and rinsed with ice-cold 1× PBS and lysed in RIPA buffer (Thermo Fisher Scientific) supplemented with 1× Halt protease and phosphatase inhibitors (Thermo Fisher Scientific). Lysis was carried out for 30 minutes at 4 °C with gentle agitation. Protein concentrations were determined using the Pierce™ Rapid Gold BCA Protein Assay Kit (Thermo Fisher Scientific). Equal amounts of protein (30 µg) were resolved by SDS-PAGE on 4-20% gradient gels and subsequently transferred onto nitrocellulose membranes. Membranes were blocked in 5% non-fat milk prepared in TBST (0.1% Tween-20 in 20mM Tris and 150mM NaCl) for a minimum of 30 minutes to reduce non-specific binding. After washing, membranes were incubated overnight at 4 °C with primary antibodies diluted in 2% non-fat milk in TBST. Primary antibodies included anti-PTEN, anti-NLRP3, anti-Akt, anti-phospho-Akt, anti-β-actin, anti-Cyclophilin B, and anti-MDH2 (Cell Signaling Technology), as well as anti-MDH1, anti-UGP2, anti-NAMPT, anti-PTPN6, and anti-INPP5D (Proteintech). All antibodies were raised in rabbits. Anti-rabbit/mouse StarBright Blue 700 secondary antibodies (Bio-Rad Laboratories) were used for detection. Fluorescent blots were directly imaged using a ChemiDoc MP imaging system (Bio-Rad Laboratories). Where appropriate, band intensities were quantified using ImageJ software.

### CHX chase assays

CHX chase assays were performed to assess the stability of candidate substrates identified in the diGly ubiquitylome analysis. BMDM^h^s (scramble control, GID8^-/-^, and MAEA^-/-^) were seeded in either 6-well plates or T-25 culture flasks prior to treatment with 5 µg/ml CHX (Cell Signaling Technology) to inhibit de novo protein synthesis and harvested at the indicated time points (0-24 hours). At each time point, cells were harvested by scraping, washed once with ice-cold PBS, and lysed directly in RIPA buffer (Thermo Fisher Scientific) supplemented with 1× Halt protease and phosphatase inhibitors, as described in the previous section. Membranes were probed with primary antibodies against NLRP3, NAMPT, MDH1, MDH2, UGP2, PTPN6, INPP5D, PTEN, or β-actin/Cyclophilin B as loading controls. Where necessary, band intensities were quantified using ImageJ software and normalized to β-actin or Cyclophilin B. For proteasome inhibition experiments, cells were treated with either CHX (5 µg/ml) or the proteasome inhibitor MG132 (10 µM) for 18 hours prior to lysis and immunoblot analysis, as described above.

### Quantification of intracellular Mtb growth in genetically modified or chemically treated BMDM^h^s

Confluent BMDM^h^ monolayers were infected with Mtb-Lux at MOI 0.5 for bioluminescence readouts or with wild-type Mtb Erdman for CFU enumeration. Growth kinetics of Mtb-Lux in infected BMDM^h^s were quantified using an EnVision plate reader (PerkinElmer). For CFU enumeration, BMDM^h^s were infected with Mtb Erdman at MOI 0.4. Following a 3-4-hour infection period to allow bacterial uptake, cells were washed at least three times with fresh macrophage culture medium to remove extracellular bacteria. At the specified time points, macrophages were lysed in sterile water containing 0.01% SDS for 15 minutes to release intracellular bacteria. Lysates were then serially diluted in 1× PBS supplemented with 0.05% Tween-80 and plated onto Middlebrook 7H10 agar supplemented with OADC. Plates were incubated at 37 °C, and CFUs were enumerated after 3-4 weeks of growth. In both cases, where applicable, compound treatments were initiated 3-4 hours post-infection (3AC and isoniazid [INH], both purchased from MedChemExpress, at final concentrations of 10 µM and 0.4 µg/ml, respectively).

### Transcriptional analysis of Mtb-infected BMDM^h^s

Scramble control and INPP5D^-/-^ BMDM^h^s were infected with the smycʹ::mCherry Mtb strain at an MOI 0.4 and were treated with 3AC beginning 3-4 hours post-infection or left untreated for 4 days, after which infected cells were flow-sorted based on mCherry expression. Sample preparation and RNA extraction were performed as previously described [25, 41], with minor modifications. Briefly, samples were treated with RNase-free Invitrogen DNase. Library preparation was carried out using Illumina’s Stranded Total RNA Prep Ligation with Ribo-Zero Plus kit and 10-bp unique dual indices. Sequencing was performed on a NovaSeq X Plus platform, generating paired-end 150-bp reads. Demultiplexing, quality control, and adapter trimming were performed using bcl-convert (v4.2.4). Due to a change in the rRNA depletion method that did not adequately remove bacterial rRNA, downstream analyses focused exclusively on the host transcriptome using analysis pipelines similar to those previously described [25, 41].

### Pathways enrichment analysis

Gene ontology (GO) pathway enrichment analysis for both proteomic and transcriptomic targets was performed in R using the enrichGO function implemented in clusterProfiler (v4.18.4) [27]. Enrichment analyses were conducted separately for Biological Process (BP), Cellular Component (CC), and Molecular Function (MF) categories. Terms were considered significantly enriched if they met a multiple testing-adjusted p-value threshold of < 0.05. For selected comparisons, the most significantly enriched GO terms (usually BPs) were further organized and visualized using bubble plots in R.

## Acknowledgements

This work was supported by funding from the National Institutes of Health (AI155319, AI162598, and OD032135), the Bill & Melinda Gates Foundation, and the Mueller Health Foundation awarded to D.G.R. Additional support was provided through a postdoctoral seed grant from the Cornell Center for Antimicrobial Resistance Research and Education to N.V.S.

## Datasets

The proteomics datasets described in this article have been deposited at the ProteomeXchange Consortium and are accessible though the accession number PXD075792. The RNA-seq datasets have been deposited at GEO Database Repository of NCBI and has an accession number of GSE325061.

